## Supplementary tables for "A computationally informed comparison between the strategies of rodents and humans in visual object recognition"

### Supplementary File 1

**Supplementary File 1a Results of binomial test on the six test protocols with the pooled data of all animals together, on the old trials (upper table) and new trials (lower table).** Here we test whether the average performance of all animals together differs significantly from chance level (indicated by p-value (50%)) or 80% (indicated by p-value (80%)) on both the old trials (upper table) as well as new trials (lower table).

| Test protocol | Mean % (old trials) | p-value (80%) | 95% CI |
| --- | --- | --- | --- |
| Rotation (x-axis) | 79.39% | 0.50 | [0.78;0.81] |
| Rotation (y-axis) | 75.86% | < .0001 | [0.74;0.78] |
| Rotation (z-axis) | 79.85% | 0.87 | [0.78;0.82] |
| Light location | 79.56% | 0.74 | [0.77;082] |
| Size | 82.73% | 0.13 | [0.79;0.86] |
| Position | 76.44% | < .0001 | [0.74;0.79] |

| Test protocol | Mean % (new trials) | p-value (80%?) | p-value (50%?) | 95% CI |
| --- | --- | --- | --- | --- |
| Rotation (x-axis) | 56.49% | < 0.0001 | < 0.0001 | [0.55;0.58] |
| Rotation (y-axis) | 66.49% | < .0001 | < 0.0001 | [0.65;0.68] |
| Rotation (z-axis) | 63.33% | < 0.0001 | < 0.0001 | [0.62;0.65] |
| Light location | 72.95% | < 0.0001 | < 0.0001 | [0.71;0.75] |
| Size | 63.49% | < .0001 | < 0.0001 | [0.57;0.64] |
| Position | 64.65% | < .0001 | < 0.0001 | [0.63;0.67] |

**Supplementary File 1b Marginal means and standard deviation of the Rotation X and Rotation Z protocols.**

| Rotation X | Target | 56.06 ± 8.54 | 56.60 ± 3.73 | 59.70 ± 5.01 | 55.66 ± 6.87 | 55.53 ± 9.54 | 55.37 ± 5.71 |
| --- | --- | --- | --- | --- | --- | --- | --- |
|  | Distractor | 64.98 ± 2.12 | 59.21 ± 6.15 | 55.07 ± 5.64 | 52.05 ± 5.64 | 56.04 ± 5.81 | 51.55 ± 4.35 |
| Rotation Z | Target | 65.40 ± 9.54 | 63.42 ± 6.03 | 62.88 ± 10.66 | 60.20 ± 8.48 | 62.58 ± 6.78 | 65.52 ± 6.91 |
|  | Distractor | 72.69 ± 4.47 | 68.25 ± 3.97 | 65.31 ± 6.27 | 60.27 ± 5.65 | 57.83 ± 6.83 | 55.64 ± 4.38 |

**Supplementary File 1c Correlation between neighbouring layers of the deep neural network.**

| Layers | Correlation |
| --- | --- |
| 1-2 | 0.11 |
| 2-3 | -0.13 |
| 3-4 | 0.05 |
| 4-5 | 0.09 |
| 5-6 | 0.12 |
| 6-7 | 0.04 |
| 7-8 | 0.17 |
| 8-9 | 0.06 |
| 9-10 | 0.01 |
| 10-11 | 0.003 |
| 11-12 | 0.11 |
| 12-13 | 0.12 |

**Supplementary File 1d Results of the linear regression model with rat performances.** A linear regression was calculated with the Classification Scores of the 13 layers as predictors and the rat performances as response vector. Each row indicates one predictor, i.e. one layer, the middle and right column indicate the t-statistic and p-value of the associated predictor, respectively.

| Predictor | T statistic | p-value |
| --- | --- | --- |
| (Intercept) | 82.15 | < 0.0001 |
| Layer 1 | 1.19 | 0.23 |
| Layer 2 | 0.56 | 0.57 |
| Layer 3 | 0.91 | 0.37 |
| Layer 4 | 0.52 | 0.60 |
| Layer 5 | -0.28 | 0.78 |
| Layer 6 | 1.38 | 0.17 |
| Layer 7 | 0.59 | 0.55 |
| Layer 8 | 2.26 | 0.025 |
| Layer 9 | 2.00 | 0.047 |
| Layer 10 | 2.35 | 0.019 |
| Layer 11 | -0.46 | 0.64 |
| Layer 12 | -0.82 | 0.41 |
| Layer 13 | 1.38 | 0.17 |

**Supplementary File 1e Results of the linear regression model with human performances.** A linear regression was calculated with the Classification Scores of the 13 layers as predictors and the human performances as response vector. Each row indicates one predictor, i.e. one layer, the middle and right column indicate the t-statistic and p-value of the associated predictor, respectively.

| Predictor | T statistic | p-value |
| --- | --- | --- |
| (Intercept) | 145.94 | < 0.0001 |
| Layer 1 | 0.63 | 0.53 |
| Layer 2 | -1.82 | 0.07 |
| Layer 3 | 0.61 | 0.54 |
| Layer 4 | 1.40 | 0.16 |
| Layer 5 | 1.61 | 0.11 |
| Layer 6 | -1.23 | 0.22 |
| Layer 7 | 0.14 | 0.89 |
| Layer 8 | 0.62 | 0.54 |
| Layer 9 | 0.80 | 0.43 |
| Layer 10 | 1.18 | 0.24 |
| Layer 11 | 2.90 | < 0.01 |
| Layer 12 | 5.08 | < 0.0001 |
| Layer 13 | 4.72 | < 0.0001 |

**Supplementary File 1f Average number of trials (SD) per test protocol and per stimulus pair (SP).** In the second column, the average number of trials per stimulus pair, for all rats together, are indicated with the standard deviation between brackets. The third column shows the total number of trials per test protocol. The fourth and fifth columns indicate the same data but for the human participants.

|  | Rats | | Humans | |
| --- | --- | --- | --- | --- |
| Protocol | Ø # trials per SP (SD) | Total # trials | Ø # trials per SP (SD) | Total # trials |
| Rotation X | 87.50 (± 9.66) | 3150 | 111.11 (± 2.55) | 4000 |
| Rotation Y | 88.22 (± 8.90) | 3176 | 111.11 (± 2.44) | 4000 |
| Rotation Z | 88.25 (± 9.55) | 3177 | 111.11 (± 3.01) | 4000 |
| Light location | 94.00 (± 11.35) | 1504 | 125 (± 3.72) | 2000 |
| Size | 205.75 (± 194.86) | 823 | 250 (± 3.16) | 1000 |
| Position | 91.28 (± 9.06) | 2282 | 120 (± 3.45) | 3000 |
| Combination rotation | 122.81 (± 12.06) | 4421 | 111.11 (± 3.45) | 4000 |
| Zero vs. high | 95.53 (± 8.76) | 4681 | 122.45 (± 4.07) | 6000 |
| High vs. zero | 109.92 (± 10.85) | 5386 | 122.45 (± 4.00) | 6000 |

**Supplementary File 1g The two criteria of choosing a CNN-informed stimulus set.** We calculated the standard deviation for the early layers and the higher layers. Based on their standard deviation, we then filtered the data and ensured that we had enough pairs per criterion.

| Criterion | Early layers | Higher layers | # pairs found |
| --- | --- | --- | --- |
| 1 | Close to zero  [-0.4*sd … 0.4*sd] | Very high  > 3*sd | 22 |
| 2 | Very high  > 3*sd | Close to zero  [-0.4*sd … 0.4*sd] | 26 |

**Supplementary File 1h Overview of the human experiment.** The left table indicates the number of trials for each training protocol. Note that for the first two phases the participants stayed in these protocols until they reached a performance of 80% or higher on the last 20 trials. The number of trials in the testing protocol is proportional to the number of pairs tested per protocol.

| Training protocol | # trials |  | Testing protocol | # trials |
| --- | --- | --- | --- | --- |
| Shaping | 80% performance or higher in the last 20 trials |  | Size | 20 |
| Training | 80% performance or higher in the last 40 trials |  | Position | 60 |
| Dimension learning | 80 |  | Light location | 40 |
| Transformations | 100 |  | Rotation X | 80 |
| Dimension learning (extra) | 20 |  | Rotation Y | 80 |
|  |  |  | Rotation Z | 80 |
|  |  |  | Combination rotation | 80 |
|  |  |  | Zero vs. high | 120 |
|  |  |  | High vs. zero | 120 |

**Supplementary File 1i An overview of the performance of the animals on the first six test protocols.** The second column indicates the planned number of sessions per animal, and between brackets the average number of sessions per animal. We excluded some sessions if the performance on the old stimuli of that session was too low. The third and fourth column provides an overview of the average performance on the old and new pairs, respectively, with the total number of trials for all rats together between brackets. We excluded the correction trials in the analysis of the old pairs. For the new pairs, no correction trials were given and reward in 80% of the trials was given.

| Test protocol | Planned # of sessions / animal (average # sessions / animal) | Mean performance on old stimuli (# old trials) | Mean performance on all new trials (# new trials) |
| --- | --- | --- | --- |
| Rotation (x-axis) | 4 (3.50) | 79.39% (1922) | 56.49% (3083) |
| Rotation (y-axis) | 4 (3.92) | 75.86% (2138) | 66.49% (3246) |
| Rotation (z-axis) | 4 (3.83) | 79.85% (2092) | 63.33% (3315) |
| Light location | 2 (1.92) | 79.56% (1043) | 72.95% (1645) |
| Size | 1 (1.00) | 82.73% (546) | 63.49% (895) |
| Position | 3 (2.83) | 76.44% (1593) | 64.65% (2423) |
