## Supplementary information for "A computationally informed comparison between the strategies of rodents and humans in visual object recognition"

### Supplementary File 2

#### Information about the pilot study

The goal of the pilot was to adjust the range of the two dimensions (concavity and alignment) in order to assure that the animals would use the two dimensions. Using the large stimulus set described in the previous paragraph, we tested a subset of them in a behavioural categorization experiment with two rats. This pilot study consisted of seven phases (see Figure 1-figure supplement 3 for an overview). We started by training rats in a base pair. This pair consists of a target and distractor that are maximally different in terms of concavity and alignment (similar as in Figure 1a), and thus are placed at the very corners of the 4x11 stimulus grid. After *Training*, we tested them in a *Dimension learning* protocol, to investigate whether the animals use both dimensions (concavity and alignment) in this discrimination task. A total of two pairs were presented to the animals. One pair consisted of the original, base pair, whereas the other pair consisted of stimuli where the alignment dimension was the opposite as in the base pair. In the third phase, *Push towards concavity,* we pushed the animals to learn the concavity dimension. A total of three pairs were presented to the animals. One pair consisted of the original training pair, whereas for the other two stimulus pairs the target and distractor differed in only one of the two dimensions, that is either concavity or alignment. The fourth pilot phase (*Push towards concavity, only new)* consisted of only the two new pairs where the target and distractor differ in only one dimension. A next phase (*More pronounced dimensions)* included stimuli where the concavity dimension was more pronounced than the base pair we started with. We changed the concavity parameter, and again the target and distractor differed in only one dimension. The sixth phase was identical to the *Dimension learning* phase, i.e. checking whether the animals use both dimensions, but this time we used the stimuli where the concavity dimension was more pronounced. In the final phase, we decided to show all four possible combinations of the stimuli.
